# Sleep Need, the Key Regulator of Sleep Homeostasis, Is Indicated and Controlled by Phosphorylation of Threonine 221 in Salt Inducible Kinase 3

**DOI:** 10.1101/2021.11.06.467421

**Authors:** Yang Li, Enxing Zhou, Yuxiang Liu, Jianjun Yu, Jingqun Yang, Chengang Li, Yufeng Cui, Tao Wang, Chaoyi Li, Ziyi Liu, Yi Rao

## Abstract

Sleep need drives sleep and plays a key role in homeostatic regulation of sleep. So far sleep need can only be inferred by animal behaviors and indicated by electroencephalography (EEG). Here we report that threonine 221 (T221) of the salt inducible kinase 3 (SIK3) was important for the catalytic activity and stability of SIK3. T221 phosphorylation in the mouse brain indicates sleep need: more sleep resulting in less phosphorylation and less sleep more phosphorylation during daily sleep/wake cycle and after sleep deprivation (SD). Sleep need was reduced in SIK3 loss of function (LOF) mutants and by T221 mutation to alanine (T221A). Sleep rebound after SD was also decreased in SIK3 LOF and T221A mutant mice. Other kinases such as SIK1 and SIK2 or other sites in SIK3 do not fulfil criteria to be both an indicator and a controller of sleep need. Our results reveal SIK3 T221 phosphorylation as the first and only chemical modification which indicates and controls sleep need.

---

Sleep is controlled by two processes: circadian (Process C) and homeostatic (Process S) ^1,2^. Process C is the circadian biological clock which has been well studied at the molecular level since the discovery of the *Drosophila Period* (*Per*) gene in 1971 ^3^ leading to the understanding of a transcriptional-translational feedback loop conserved from insects to mammals including humans ^4-9^. By contrast, our understanding of Process S, the key component of homeostatic regulation of sleep, lags far behind that of Process C.

Process S is regulated by sleep need ^1,2,10,11^, which can be measured by the spectral power in the δ range (1-4 Hz) electroencephalogram (EEG) of non-rapid eye movement sleep (NREMS) ^11-14^.

Sleep need is regulated by prior wakeful experience ^1,15^. During a natural day, there is gradual loss of sleep and concomitant increase of sleep need due to daily activities (in the light phase for humans or the dark phase for mice), and a gradual decrease of sleep need due to sleep (in the dark phase for humans or the light phase for mice). Sleep need can also be increased by artificial sleep deprivation (SD): after the natural sleep is deprived (during the dark phase of humans or the light phase of mice), there is an increase of sleep need. Sleep is homeostatically regulated: loss of sleep in a natural day will be followed by sleep and artificial SD will result in a rebound of sleep.

A controller for sleep need should be necessary for EEG delta power in NREMS before and after SD, as well as sleep rebound after SD. So far, no molecule or chemical modification has been shown to be functionally required for all three (EEG δ power before SD, EEG δ power after SD and sleep rebound after SD).

An ideal indicator for sleep need should show correlation with sleep need in a natural sleep/wake cycle as well as that upon SD. So far, no molecule or chemical modification fulfil both criteria of a sleep need indicator.

In flies, we and others have found multiple genes whose loss of function (LOF) mutations significantly decreased sleep rebound after SD ^16-26^. However, there is no EEG delta power equivalent in flies to measure sleep need. No molecule or chemical modification could be established as an indicator of sleep need in flies.

In mice, the salt inducible kinase 3 (SIK3) has been implicated in sleep regulation ^27^. It is based on the finding of a dominant gain of function (GOF) mutation *Sleepy* which eliminated a segment containing serine 551 (S551), a target site for protein kinase A (PKA). Removal of S551 was known to increase SIK3 activity ^28^. *Sleepy* increased sleep need ^27^. Removal of S551 was known to increase SIK3 activity ^28^. S551 mutation to alanine (A) (S551A) increased sleep need ^29^. However, both Sleepy and S551A are GOF mutations with increased SIK3 activities ^28^. LOF SIK3 mutations have not been studied and therefore physiological requirement of SIK3 in sleep need could not be established.

There are three SIKs in mammals ^30-33^. Mutations in SIK1 (S577A) and SIK2 (S587A) are equivalent to S551A in SIK3 and they also increased sleep need in mice ^34^. Thus, all three SIKs were implicated in sleep need by GOF mutations, but none of the wild type (wt) mouse SIK genes has been shown to be physiologically necessary for sleep need, which requires evidence from LOF mutations. As we will show here, LOF SIK1 and SIK2 mutants were similar to the wt in sleep need, thus, neither SIK1 nor SIK2 is necessary for sleep need. It is essential to investigate LOF SIK3 mutants to determine whether SIK3 is necessary for sleep need.

While S551A of SIK3 increased sleep need ^29^, S551 phosphorylation did not change after SD ^27,29^. Thus, S551 phosphorylation does not indicate sleep need. S577 phosphorylation in SIK1 and S587 phosphorylation in SIK2 have not been previously studied *in vivo*, but it may not be productive to examine the phosphorylation of either SIK1 S577 or SIK2 S587 in light of our findings that neither SIK1 nor SIK2 is necessary for sleep need in mice.

Here we show T221 phosphorylation of SIK3 satisfies all criteria for a sleep need indicator and those for a sleep need controller.

## RESULTS

### Specific Recognition of Phosphorylated T221 in SIK3 by a Monoclonal Antibody *in vitro* and *in vivo*

SIKs from different species share a general structure with a catalytic domain, an ubiquitin-associated domain and a C terminal domain ^35-37^. There are multiple sites of phosphorylation in a SIK protein, among which targets for PKA have been well studied: PKA usually negatively regulates SIK activities ^28,38-42^.

T221 of SIK3 is not a target site for PKA. It was found to change after SD by mass spectrometry (MS) ^27^, but it has not been studied whether its level changes during a natural day of sleep/wake cycle. In order to examine natural changes more easily than relying on MS, we needed an antibody which could recognize phosphorylated T221. We tested the specificity of a rabbit monoclonal antibody (MAb) and found that it could recognize the phosphorylated T221 of SIK3 from human embryonic kidney (HEK) cells transfected with a cDNA expressing SIK3 fused inframe to the FLAG epitope tag (Fig. 1a). The MAb did not recognize T221 of SIK3 if the cell lysate was treated with phosphatase (Fig. 1a). The MAb also did not recognize SIK3 if the amino acid residue at 221 was mutated to A (T221A) or to glutamic acid (T221E) in HEK cells (Fig. 1a). These results show that T221 phosphorylated in HEK cells is recognized by the anti-SIK3 phospho-T221 MAb.

**Figure 1.**
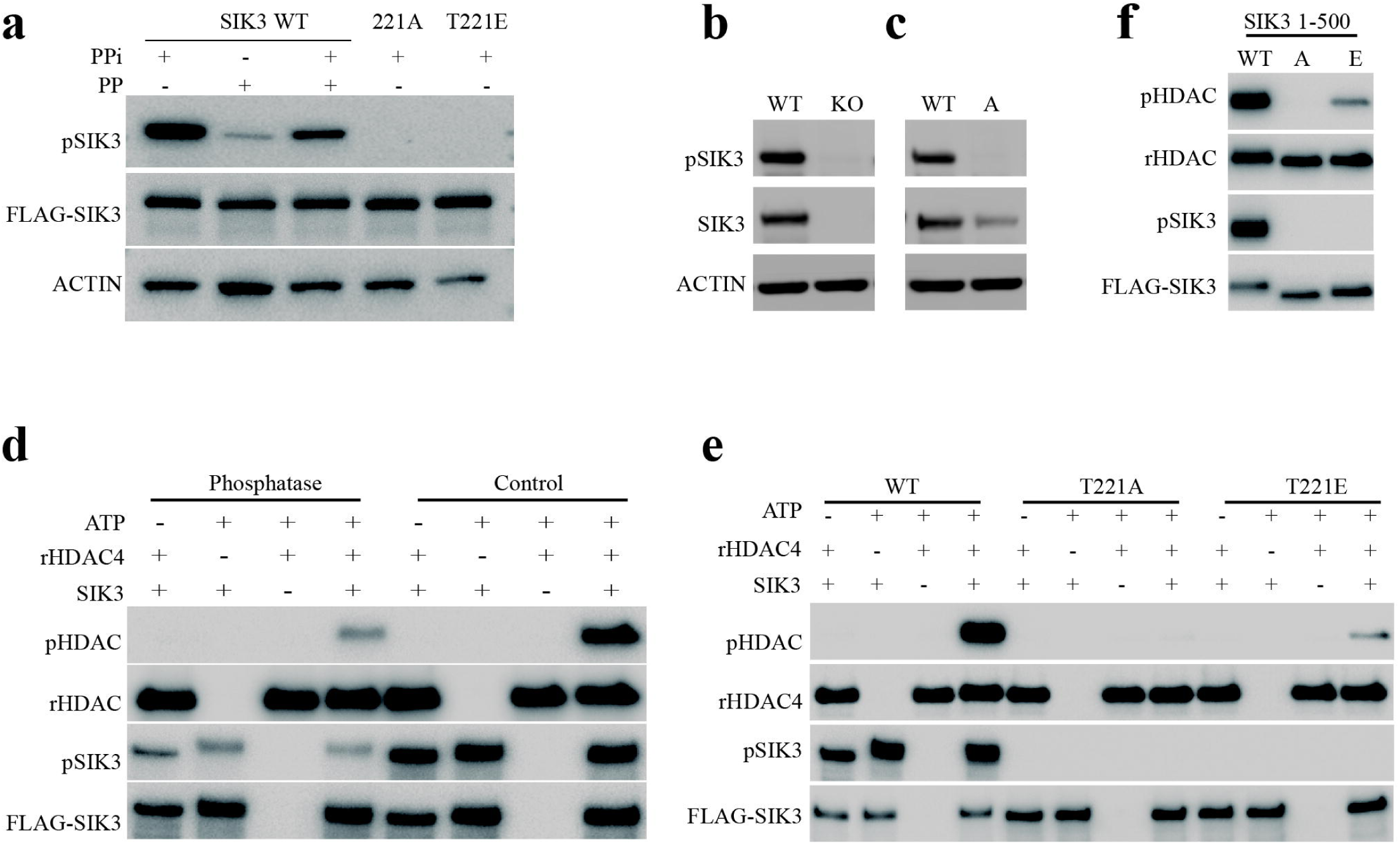
T221 Phosphorylation and SIK3 Catalytic Activity. **a**. Specificity of anti-SIK3 pT221 mAb in HEK293T cells. HEK293T cells were transfected with indicated plasmids and harvested 24 hrs after transfection. Extracts were subjected to Western blot analysis (WB) with indicated antibodies. (“p” indicates phosphosphorylated amino acid residues) **b**. Immunoblotting with anti-SIK3 pT221 and anti-SIK3 antibodies of brain homogenates from *Sik3*^*+/+*^ and *Sik3*^*-/-*^ mice. **c**. Immunoblotting with anti-SIK3 pT221 and anti-SIK3 antibodies of brain homogenates from *Sik3* ^*+/+*^ and *Sik3*^*221A/A*^ mice. **d**. SIK3 is regulated by phosphorylation at T221 site. FLAG-SIK3 was immunoprecipitated and pre-treated with or without λ-phosphatase for 1 hr at 37□. HDAC4 phosphorylation was detected by an antibody against phospho-HDAC4. **e**. The enzymatic activity of SIK3 WT protein is higher than those of T221E and T221A SIK mutant proteins. Indicated proteins were immunoprecipitated and assayed as in d. **f**. The enzymatic activity of SIK3 1-500 WT protein is er than those of T221E and T221A SIK mutant proteins. Indicated proteins were immunoprecipitated and assayed as in d.

To examine whether T221 phosphorylated in mice can be specifically recognized by the anti-SIK3 phospho-T221 MAb, we eliminated the SIK3 gene by gene targeting. We found that SIK3 knockout caused embryonic lethality, with a few mice born with the SIK3^-/-^ genotype among hundreds of wt mice. While the number of SIK3^-/-^ mice were too low to allow behavior or EEG analysis, we did manage to use them for protein analysis to confirm that the protein recognized by the MAb was absent in SIK3^-/-^mice while that in the wt was recognized by both the anti-SIK3 antibodies and the anti-SIK3 phospho-T221 MAb (Fig. 1b).

By the CRISPR-CAS9 method, we generated mice carrying the point mutation T221A (Fig. 5). SIK3 protein was detected in T221A mutant mice, whereas T221A was not recognized by the MAb (Fig. 1c), again demonstrating the specificity of the anti-phospho-T221 MAb in mice.

Taken together, our results provide *in vitro* and *in vivo* evidence that phospho-T221 in SIK3 can be recognized specifically by the anti-SIK3 phospho-T221 MAb.

### Enhancement of SIK3 Catalytic Activity by T221 Phosphorylation

Histone deacetylase 4 (HDAC4) is a known substrate of SIK3 ^43^. We used it to test the catalytic activities of SIK3 and its mutants. We transfected HEK cells with a cDNA construct expressing the FLAG-tagged SIK3 and found SIK3 immunoprecipitated with the anti-FLAG antibody to be active in phosphorylating HDAC4 (Fig. 1d “Control” lanes on the right), but this activity was reduced if SIK3 was treated with phosphatase (Fig. 1d “Phosphatase” lanes on the left).

T221A mutant of the full length SIK3 expressed in HEK cells was not active in phosphorylating HDAC4 (Fig. 1e).

A shorter version of SIK3 containing amino acid residues 1 to 500 was also active in phosphorylating HDAC4 (Fig. 1f, WT), but T221A mutant of SIK3 1-500 was inactive (Fig.1f, a). These results show that T221 phosphorylation enhances SIK3 catalytic activity.

Surprisingly, the activity of SIK3 T221E immunoprecipitated from HEK was lower than the wt SIK3 in phosphorylating HDAC4 (Fig. 1e), though T221E was higher than T221A in phosphorylating HDAC4 (Fig. 1e). T221E mutation in SIK3 1-500 reduced but did not eliminate its activity (Fig. 1f).

Results from SIK3 catalytic activity show that SIK3 T221A could be used as an LOF mutant for SIK3, whereas SIK3 T221E could not be used as a GOF mutant for SIK3. We note that, rather than being more active mutants, S to D mutations in SIK3 (S551D), SIK1 (S587D) and SIK2 (S577D) were all less active than their corresponding wt ^29,34^ with S to D mutants ending up with phenotypes qualitatively similar to those of S to A mutants ^29,34^. In the case of these SIK phospho-site mutants, it was not helpful to use E or D mutants to find mutants more active than wt SIK proteins, making it meaningful to examine the *in vivo* phenotypes of only S or T to A mutants.

### Enhancement of SIK3 Protein Stability by T221 Phosphorylation

We obtained *Sik3 knockout first* (*Sik3-kof*) mutant mice into whose *Sik3* gene the bacterial *lacZ* gene was inserted (Fig. 2a). *Sik3-kof*/+ and *Sik3-kof*/*Sik3-kof* were both viable with dosage-dependently reduction of SIK3 mRNA expression (Fig. 2b) and SIK3 protein (Fig. 2d). The anti-phospho-T221 MAb could recognize a band in *Sik3-kof*/+ and *Sik3-kof*/*Sik3-kof* mice but the level of phospho-T221 also depended on the gene dosage (Fig. 2d). Therefore, *Sik3-kof* was used as a partial LOF mutant for SIK3.

**Figure 2.**
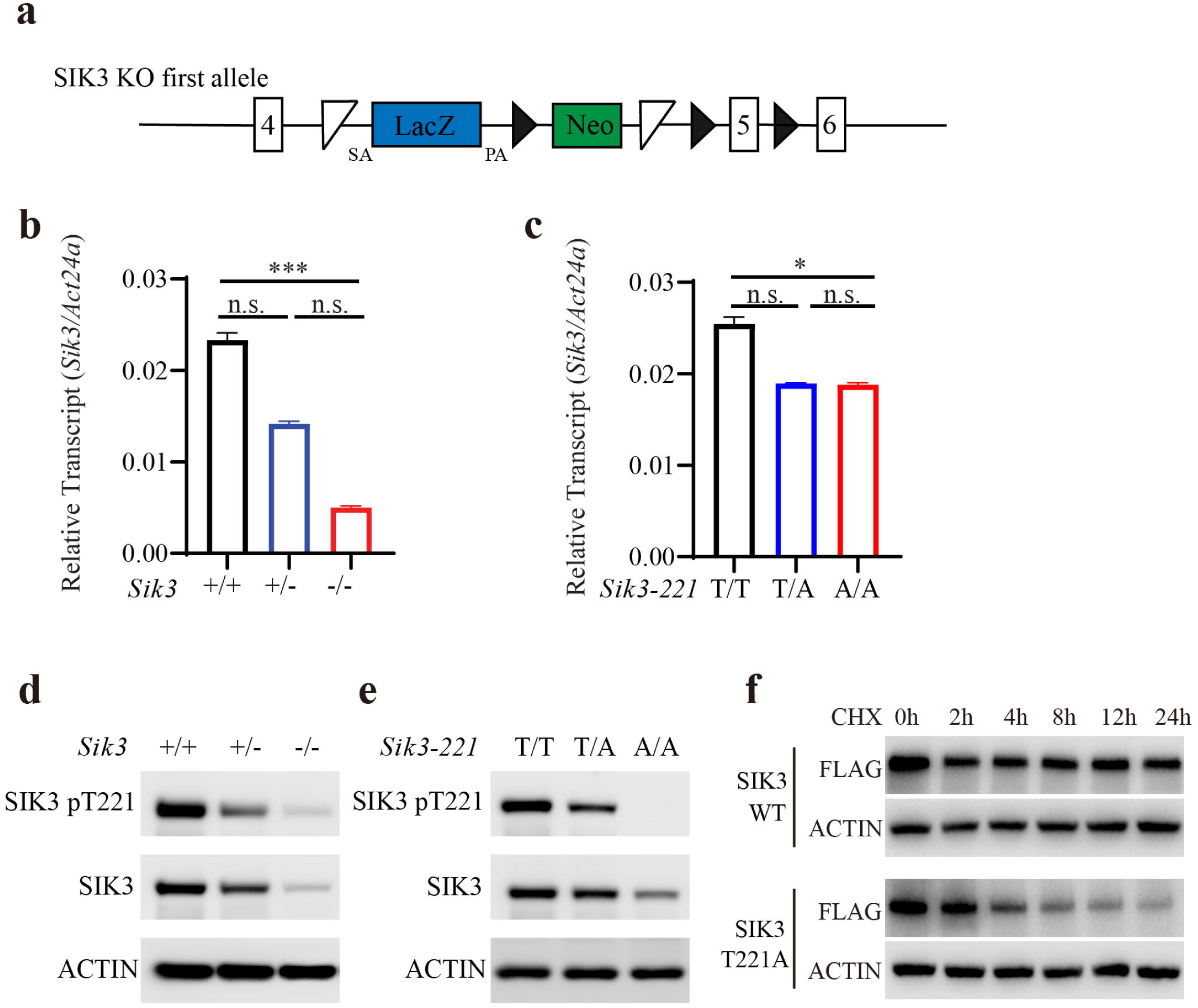
Dynamics of SIK3 Protein and T221 Phosphorylation in a Sleep/Wake Cycle and after Sleep Deprivation. **a**. Temporal profile of SIK3 protein level and T221 phosphorylation level in mouse brains (n=12 at each time point). PER2 protein level and PER2 S662 phosphorylation level are also shown. ZT: zeitgeber time **b-h**. The effect of sleep deprivation on SIK3 and T221 phosphorylation in mouse brains. Immunoblotting with anti-SIK3 T221 antibody **(b)**, anti-SIK3 antibody **(c)**, anti-actin antibody **(d)**, anti-phospho-ERK1/2 antibody **(d)**, anti-ERK1/2 antibody **(f)**, anti-phospho-JNK antibody **(g)**, anti-JNK antibody **(h)**. SD: 6 hrs of SD after ZT 0.

With our T221A mutant mice, we found that the mRNA levels of SIK3 in +/221A and 221A/221A mice were similar and both were lower than that in the wt (Fig.2c). It could also recognize a band in +/221A mice but not in 221A/221A mice (Fig. 2e). All these results further confirmed the specificity of the MAb *in vivo*.

Interestingly, while the mRNA levels of SIK3 were similar in +/221A and 221A/221A mice, the level of SIK3 protein was lower in 221A/221A mice than that in +/221A mice (Fig. 2e). Phospho-T221 could be detected in +/221A mice but not in 221A/221A mice (Fig. 2e).

While the reduction of SIK3 mRNA level could partially explain the reduction of SIK3 protein in 221A/221A mice compared to the wt (+/+) (Fig. 1c, Fig. 2e), the finding that the level of SIK3 protein in 221A/221A mice was lower than that in +/221A mice could only be explained by reduced protein synthesis or stability caused by T to A mutation at residue 221 of SIK3.

Phosphorylation is well known to regulate protein stability. For example, in the case of the PER protein involved in circadian regulation of animals from *Drosophila* to humans, phosphorylation at multiple sites regulates its stability ^9,44,45^.

To directly investigate SIK3 protein stability, we transfected cDNA constructs expressing either wt SIK3 or T221 SIK3 with a FLAG tag into HEK cells. After cycloheximide treatment to inhibit protein synthesis, levels of SIK3 were followed to examine SIK3 protein stability over the next 24 hours (hrs) (Fig. 2f). There was a reduction of SIK3 protein in 2 hrs, but it was stable for the next 22 hrs (Fig. 2f). By contrast, the level of SIK3 T221A protein decreased continuously over 24 hrs (Fig. 2f).

Thus, both *in vivo* (Fig. 2e) and *in vitro* (Fig. 2f) results are consistent with the idea that T221 phosphorylation enhances the stability of SIK3 protein.

### Indication of Sleep Need by T221 Phosphorylation

Sleep need decreases with more sleep and increases with less sleep ^11-15^.To investigate whether T221 phosphorylation in SIK3 indicates sleep need, we first examined T221 phosphorylation in daily natural sleep/wake cycle. Mice were kept at a 12 hr dark/12 hr light schedule, with zeitgeber time (ZT) 0 as the beginning of the light phase when mice started to sleep and ZT12 as the beginning of the night when mice woke up.

We collected brains from 12 mice every 3 hrs over 36 hrs. While the level of actin did not change, T221 phosphorylation was highest at ZT0, decreasing to its lowest point at ZT12, returning gradually to peak again at ZT24 before decreasing again for another 12 hrs (Fig. 3a).

**Figure 3.**
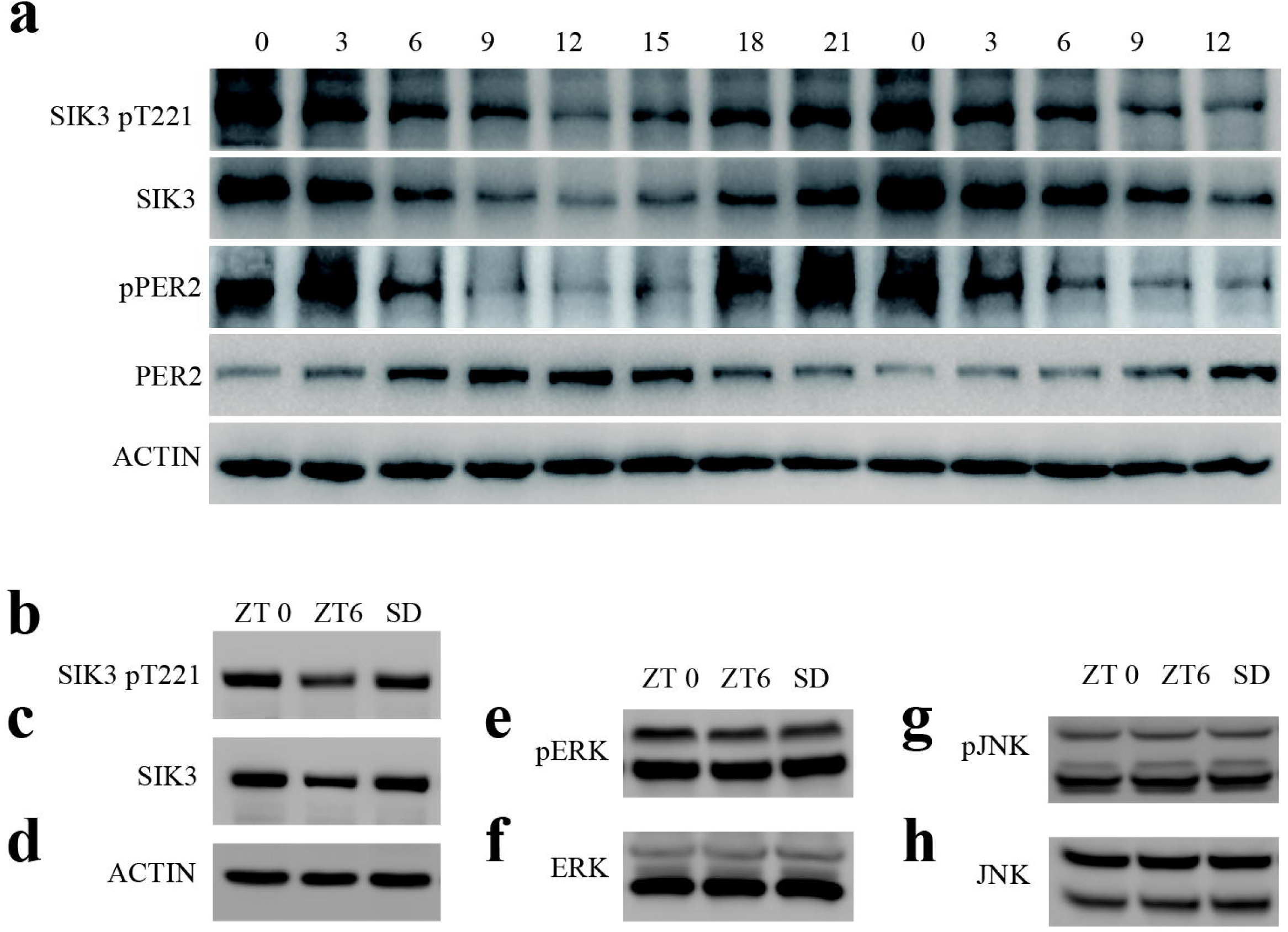
T221 Phosphorylation and SIK3 Protein Stability. **a**. A schematic diagram of the *Sik3-kof* allele. Insertion of lacZ-polyA into intron 4 leads to *Sik3* transcription termination after exon 4. The white triangle represents FRT, and the black triangle represents Loxp. SA: splice acceptor, PA: poly A. **b**. Relative *Sik3* mRNA expression levels in the brains of *Sik3*^*+/+*^(n=6), *Sik3*^*+/-*^ (n=6) and *Sik3*^*-/-*^ (n=6) mice. Kruskal-Wallis test with Dunn’s posttest. \*\*\**P<0*.*001*. Error bars represent standard erroe of the mean (SEM). **c**. Relative *Sik3* mRNA expression levels in the brains of *Sik3*^*+/+*^(n=3), *Sik3*^*+/221A*^(n=3) and *Sik3*^*221A/A*^ (n=3) mice. Kruskal-Wallis test with Dunn’s posttest. \**P<0*.*05*. Error bars represent SEM. **d**. Levels of SIK3 T221 phosphorylation and SIK3 protein expression in the brains from *Sik3*^*+/+*^, *Sik3*^*+/-*^ and *Sik3*^*-/-*^ mice. **e**. Levels of SIK3 T221 phosphorylation and SIK3 protein expression in the brains of *Sik3*^*+/+*^, *Sik3*^*+/221A*^ and *Sik3*^*221A/A*^ mice. **f**. SIK3 protein degradation analysized in HEK293T cells. Cells were transfected with SIK3 WT or T221A plasmid and cultured for 24 hrs. Transfected cells were treated with 200 mM cycloheximide (CHX) before harvest at indicated time points. Equal amounts of extracts were subjected to immunoblot analysis with indicated antibodies.

The correlation of the level of T221 phosphorylation with the level of SIK3 protein further supports our conclusion from previous studies (Fig. 2e and 2f) that T221 phosphorylation stabilizes SIK3 protein.

By contrast, the level of mouse PER2 protein increased from ZT0 to ZT12 before decreasing to reach its lowest level at ZT24 (Fig. 3a). The level of PER2 phosphorylation at S662 was opposite to the level of PER2 protein, consistent with the prior knowledge that S662 phosphorylation destabilizes PER2 protein ^45,46^.

We used the MAb recognizing phospho-T221 to confirm increased T221 phosphorylation after 6 hrs of SD (Fig. 3b), which was previously found by mass spectrometry ^27^. Consistent with our demonstration that T221 phosphorylation stabilizes SIK3 protein, the level of SIK3 protein also correlated with SIK3 phosphorylation with and without SD (Fig. 3c). The levels of kinases such as ERK and JNK did not change with SD (Figure 3f and 3h), nor the levels of ERK or JNK phosphorylation changed with SD (Fig. 3e, 3g).

Our results reveal that the phosphorylation of SIK3 at T221, but not the phosphorylation of other kinases such as ERK or JNK, indicates sleep need.

### Lack of Requirement for SIK1 and SIK2 in Sleep Need

SIK3 was first discovered for its role in sleep through *Sleepy* mutants, in which a segment in SIK3 including S551 was deleted, alleviating PKA mediated negative regulation ^27^. S551 in SIK3 are equivalent to S577 in SIK1 and S587 in SIK2 ^34^. Single point mutations of SIK3 S551A, SIK1 S577A and SIK2 S587A all led to loss of PKA phosphorylation of these sites and increased their function ^29,34^. All three point mutations increased sleep need in mice ^29,34^. While these results suggest that all three SIK proteins regulate sleep need, the very nature of GOF made it not possible to establish a physiological requirement for any of the SIK proteins in regulating sleep need.

To investigate whether SIK1 or SIK2 is required for sleep need, we generated mice with their SIK1 or SIK2 gene deleted (Extended Data Fig. 1 and 2). Unlike SIK3 knockout mutation which caused embryonic lethality, these knockout mice were viable and were subject to our experimental analysis of sleep need.

Our knockout strategies eliminated the expression of SIK1 (Extended Data Fig. 1f) and SIK2 mRNAs (Extended Data Fig. 2f).

δ power is a measure of EEG activity in the 1–4 Hz range, in slow-wave sleep (SWS, NREM sleep in humans), and reflects sleep need ^1,10-14^.

No parameters for sleep or sleep need were changed in either kind of mutants (Extended Data Fig. 1 and 2). Most clearly, sleep need as measured by δ power EEG was not significantly different among SIK1^+/+^, SIK1^+/-^ or SIK1^-/-^ mice (Extended Data Fig. 1d) at any time point over 24 hrs (Extended Data Fig. 1e). Similarly, sleep need as measured by δ power EEG was not different among SIK2^+/+^, SIK2^+/-^ or SIK2^-/-^ mice (Extended Data Fig. 2d) at any time point over 24 hrs (Extended Data Fig. 2e).

These results show that neither SIK1 nor SIK2 is physiologically required for sleep need.

### Functional Significance of SIK3 in Sleep Need

SIK3 elimination caused embryonic lethality and those that were born grew slowly, leaving us without sufficient number of SIK3 knockout mice for EEG analysis. Partial LOF *Sik3-kof* mutants (Fig. 2a, 2b, 2d) allowed us to investigate physiological requirement of SIK3 in sleep need.

Daytime wake duration was significantly longer in *Sik3-kof*/*Sik3-kof* (-/-) mice when compared with +/+ mice (Figure 4A). Daytime rapid eye movement (REM) sleep duration was significantly less in *Sik3-kof*/*Sik3-kof* (-/-) mice than that in +/+ mice (Fig. 4a). Sleep or wake episode durations and episode numbers were not significantly different between *Sik3-kof*/*Sik3-kof* (-/-) mice and wt mice (Extended Data Fig. 3a, 3b).

**Figure 4.**
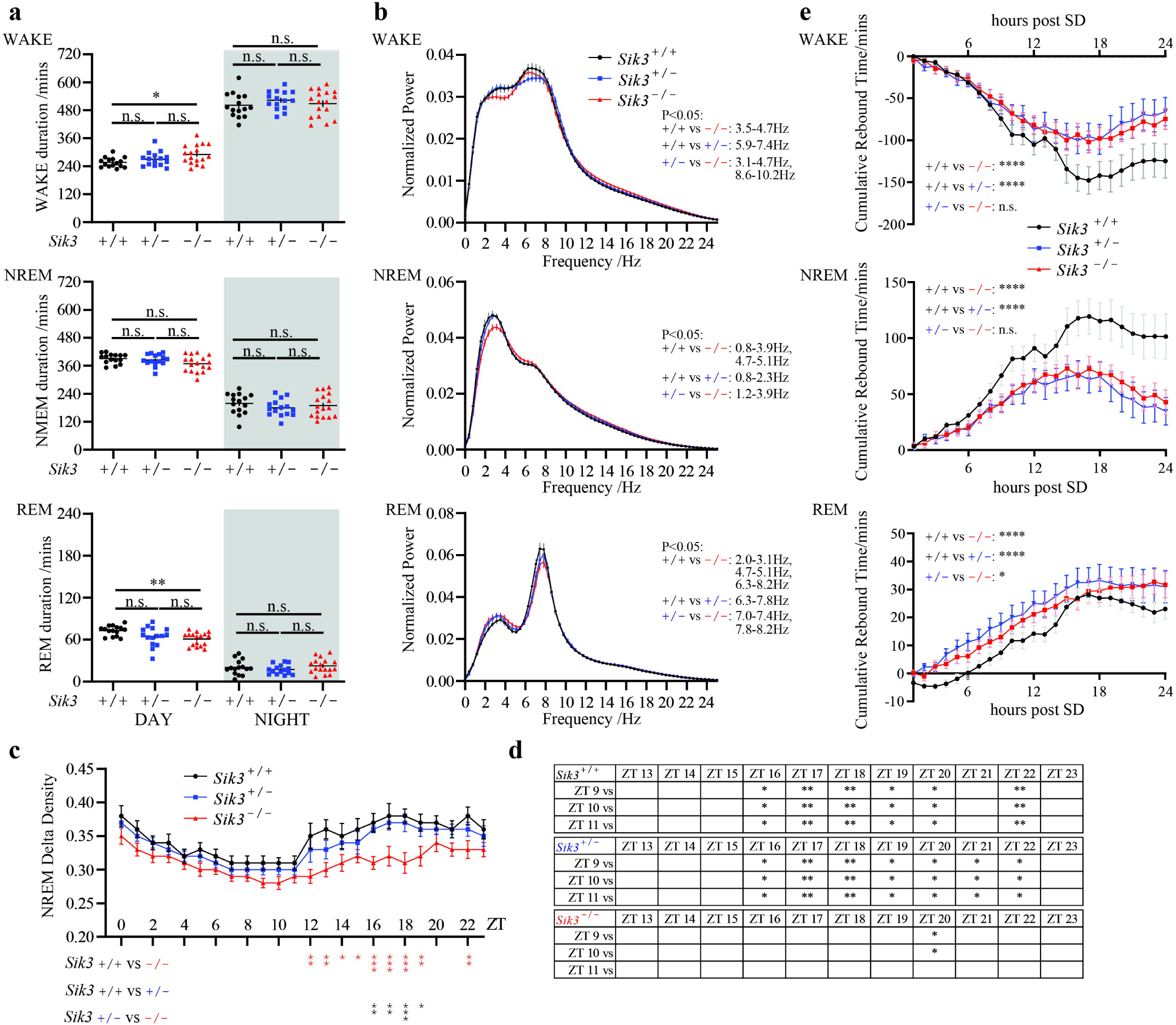
SIK3 and Sleep Need in Mice. **a**. The duration of WAKE, NREM and REM in *Sik3*^*+/+*^(n=15), *Sik3/Sik3kof* (^*+/-*^) (n=15) and *Sik3kof/Sik3kof* (^*-/-*^) (n=17) mice. \**P*< 0.05, \*\**P*<0.01, Kruskal-Wallis test with Dunn’s posttest. Error bars represent SEM. **b**. EEG power spectrum of WAKE, NREM and REM states in *Sik3*^*+/+*^(n=15), *Sik3*^*+/-*^(n=15) and *Sik3*^*-/-*^(n=17). Two-way ANOVA with Tukey’s posttest, *P*<0.05 pairs were listed on the right. Error bars represent SEM. **c-d**. NREM δ-power density of *Sik3*^*+/+*^(n=15), *Sik3*^*+/-*^(n=15) and *Sik3*^*-/-*^(n=17). Two-way ANOVA with Tukey’s posttest, statistical comparison among genotypes listed below, and statistical comparison among different times listed on **D**. \**P*<0.05, \*\**P*<0.01, \*\*\**P*<0.001. Error bars represent SEM. **e**. Accumulated WAKE loss, and NREM, REM sleep gain of *Sik3*^*+/+*^(n=11), *Sik3*^*+/-*^(n=12) and *Sik3*^*-/-*^(n=11) after 6 hrs of SD. Two-way ANOVA with Tukey’s posttest, \**P*<0.05, \*\*\*\**P*<0.0001, Error bars represent SEM.

Analysis of the EEG in *Sik3-kof* mutants showed that, during the NREM stages, power spectrum in the delta range (1-4Hz) was significantly decreased: those in *Sik3-kof*/*Sik3-kof* (-/-) mice were significantly lower than those in +/+ and +/-mice (Fig. 4b). Delta power density (the ratio of 1-4 Hz delta power to overall 1-25 Hz EEG power) during NREM over a 24 hrs plot was significantly reduced in *Sik3-kof* mutants (Fig. 4c), especially during the night phase when sleep need accumulated in mice, at multiple time points, NREM δ density in *Sik3-kof*/*Sik3-kof* (-/-) mice was significantly lower than those in +/+ and +/-mice (Fig. 4c).

Sleep need decreased gradually during light phase, when mice usually fell asleep, and increased gradually during dark phase, when mice were often awake. In the wt, δ power densities at ZT16, 17, 18, 19, 20 and 22 were all significantly higher than those at ZT9, 10 and 11 (Fig. 4d). In *Sik3-kof*/*+* (-/+) mice, δ power densities at ZT16, 17, 18, 19, 20, 21 and 22 were all significantly higher than those at ZT9, 10 and 11 (Fig. 4d). However, in *Sik3-kof*/*Sik3-kof* (-/-) mice, only δ power density at ZT20 was significantly higher than those at ZT9 and ZT10, with no significantly at any other time point (Fig. 4d). These results indicate that SIK3 was necessary for accumulation of sleep over the active phase in mice. In short, no accumulation of sleep need in the absence of SIK3.

To investigate involvement of SIK3 in homeostatic sleep regulation, we carried out 6 hrs of SD and analyzed changes of sleep homeostasis. In wt mice, NREM and REM rebounded after SD, with WAKE duration reduced (Fig. 4e). NREM rebound and WAKE reduction were significantly less in *Sik3-kof* mutant mice than those in wt mice (Fig. 4e). Curiously, REM rebounded more in *Sik3-kof* mutant mice than that in wt mice.

Results *Sik3-kof* mutant mice show that SIK3 is required for sleep need and for sleep rebound after SD.

### Functional Significance of SIK3-T221 Phosphorylation in Sleep Need

Our SIK3-T221A mutant mice (Fig. 2c and 2d) allowed investigation of the physiological requirement of T221 phosphorylation in sleep need.

T221A mutants exhibited shorter NREM sleep duration (Fig. 5a), especially during the dark phase, with no change in REM sleep duration. Sleep or wake episode durations and episode numbers were not significantly different between T221A mutant mice and wt mice (Extended Data Fig. 4a, 4b). The specific decrease of NREM sleep duration in T221A mutant mice suggests possible reduction of homeostatic sleep pressure.

**Figure 5.**
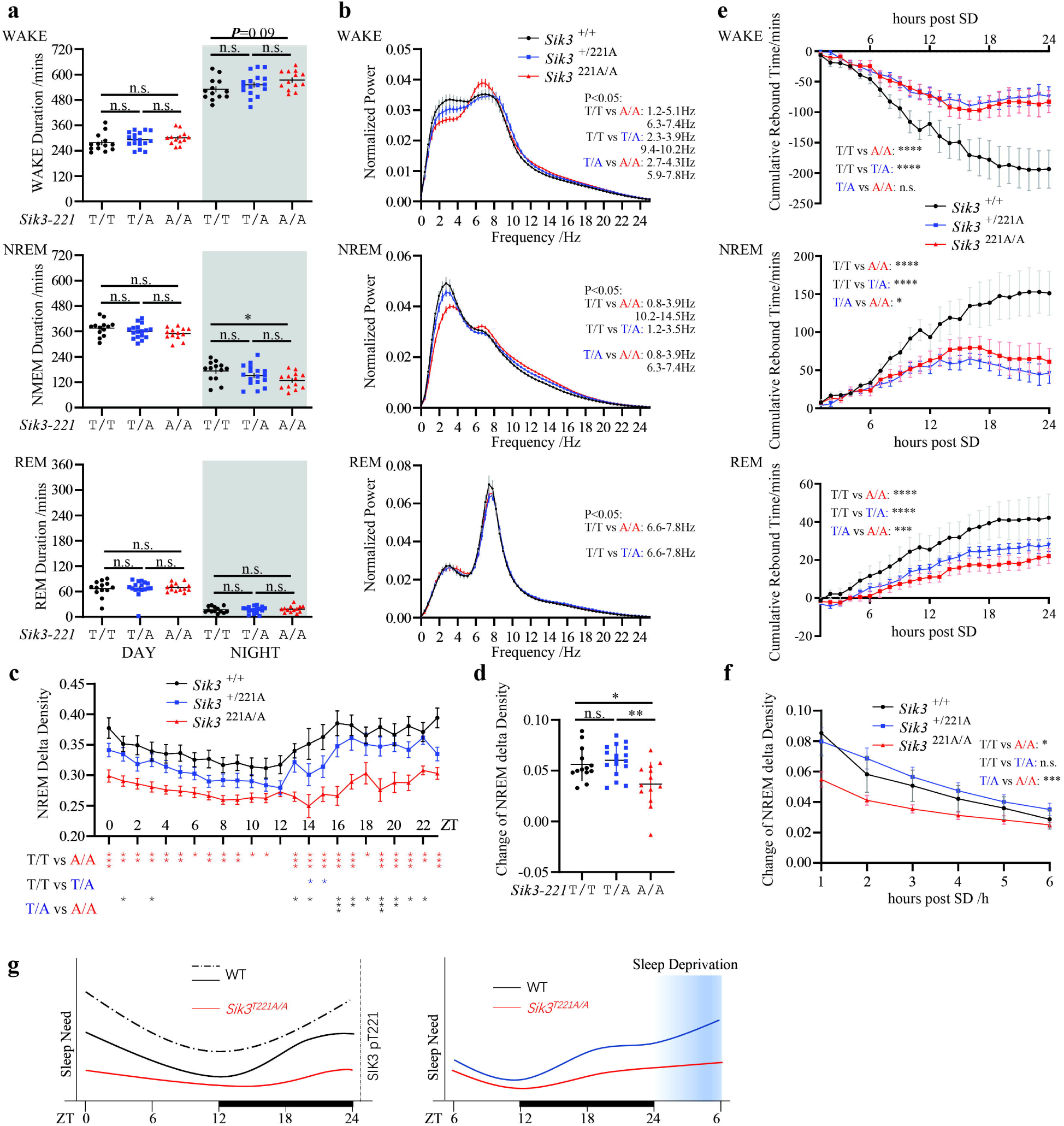
T221 Phosphorylation and Sleep Need in Mice. **a**. The duration of WAKE, NREM and REM in *Sik3*^*+/+*^(n=13), *Sik3*^*+/221A*^(n=17) and *Sik3*^*221A/221A*^(n=13). \**P*< 0.05, Kruskal–Wallis test with Dunn’s posttest. Error bars represent SEM. **b**. EEG power spectrum of WAKE, NREM and REM states in *Sik3*^*+/+*^(n=13), *Sik3*^*+/221A*^(n=17) and *Sik3*^*221A/221A*^(n=13). Two-way ANOVA with Tukey’s posttest, the *P*<0.05 pairs was listed on the right. Error bars represent SEM. **c**. NREM δ-power density of *Sik3*^*+/+*^(n=13), *Sik3*^*+/221A*^(n=17) and *Sik3*^*221A/221A*^(n=13). Two-way ANOVA with Tukey’s posttest, statistical comparison among genotypes were listed below. \**P*< 0.05, \*\**P*<0.01, \*\*\**P*<0.001. Error bars represent SEM. **d**. δ power density increasement from ZT9-11 to ZT19-24 of *Sik3*^*+/+*^(n=13), *Sik3*^*+/221A*^(n=17) and *Sik3*^*221A/221A*^(n=13). Kruskal–Wallis test with Dunn’s posttest. \**P*< 0.05, \*\**P*<0.01. Error bars represent SEM. **e**. Accumulated WAKE loss, and NREM, REM sleep gain of *Sik3*^*+/+*^(n=6), *Sik3*^*+/221A*^(n=10) and *Sik3*^*221A/221A*^(n=8) after 6 hrs SD. Two-way ANOVA with Tukey’s posttest, \**P*< 0.05, \*\*\**P*<0.001, \*\*\*\**P*<0.0001, Error bars represent SEM. **f**. δ power density changes after SD at corresponding time points of *Sik3*^*+/+*^(n=6), *Sik3*^*+/221A*^(n=10) and *Sik3*^*221A/221A*^(n=8) mice. For a given duration of time (T), changes(T)=post SD density[T]–pre SD density[T], Density[T] denotes the mean NREM δ power density within T. Two-way ANOVA with Tukey’s posttest, \**P*< 0.05, \*\*\**P*<0.001, Error bars represent SEM. **g**. A schematic illustration how SIK3-T221 phosphorylation indicates and controls sleep need in mice.

EEG spectrum analysis showed reduced δ power density during NREM in T221A/T221A mutant mice as compared to wt mice (Fig. 5b). During the light phase, T221A mutants were similar to wt mice in decreasing sleep need, as the δ power density decreased at a similar pace in T221A mutant and wt mice (Fig. 5c). During the dark phase the delta power of wt mice increased significantly whereas that in T221A mutants did not significantly change (Fig. 5c). The increase of δ power density from ZT9-12 (when mice had slept for a day) to ZT17-24 (when mice had been awake for a night) was significantly less in T221A mutant mice than that in wt mice (Fig. 5d). δ power EEG of T221A mutants were similar to wt mice during REM sleep (Fig. 5b). At all points over 24 hrs, NREM delta power of T221A mutants was significantly less than that in the wt mice (Fig. 5c), also consistent with a role for T221 in sleep need.

These results indicate that T221 phosphorylation in SIK3 was required for sleep need.

We then analyzed rebound after SD in T221A mutant mice. After 6 hrs of SD from the onset of light phase, mice began to regain their lost NREM and REM sleep stages during the following 24 hours (Fig. 5e). Both the NREM rebound and the REM rebound were significantly less in T221A mutant mice than those in wt mice (Fig. 5e).

δ density after SD was significantly lower in T221A mutant mice than that in wt mice (Extended Data Fig. 4c). Changes of δ density after SD from corresponding baseline levels were also significantly less in T221A mutant mice than those in wt mice (Fig. 5f). Over the entire 24 hrs after SD, NREM δ density was lower in T221A mice than that in the wt (Extended Data Fig. 4c).

It is notable that the sleep need phenotypes of T221A mutants (Fig. 5 and Extended Data 4) were more significant than *Sik3-kof* mutants (Fig. 4 and Extended Data Fig. 3).

These results show that T221 phosphorylation in SIK3 was required for sleep need, for changes of sleep need during a regular daily sleep/wake cycle and for changes of sleep need after SD. T221 is also required for sleep rebound after SD.

## DISCUSSION

Our results have demonstrated that phosphorylation of T221 in SIK3 is both an indicator and controller of sleep need in mammalian animals. Because other candidates, such as other kinases (SIK1 and SIK2) or the S551 in SIK3, do not satisfy the criteria, SIK3-T221 is not only the first but also the only indicator and controller of sleep need, at this point.

Our model is summarized in Fig. 5g: the left panel shows a natural daily sleep/wake cycle. The two solid lines represent sleep need (wt in black and SIK3 T221 mutants in red) and the dotted line represents T221 phosphorylation in the wt mice. Sleep need and SIK3-T221 phosphorylation increase during the WAKE stages (or the dark phase for mice); sleep need and SIK3-T221 phosphorylation decrease during the SLEEP stages (or the light phase for mice). When T221 is mutated to A221, the increase of sleep need during the WAKE stages is significantly reduced. The right panel shows effects of SD. Sleep need and T221 phosphorylation are increased by SD. T221A mutation reduces sleep need increase caused by SD and reduces sleep rebound after SD.

This model is supported by multiple pieces of evidence presented in this paper. We have found that sleep need and sleep need accumulation in mice were decreased by a LOF mutation of SIK3 (*Sik3-kof*) (Fig. 4), demonstrating a physiological requirement of SIK3 in sleep need. Phosphorylation of T221 in SIK3 monitored sleep need in mice: T221 phosphorylation increased with the loss of sleep during a natural daily sleep/wake cycle and T221 phosphorylation decreased with sleep (Fig. 3a). Artificial SD increased T221 phosphorylation (Fig. 3b). Functionally, sleep need and need accumulation rate were reduced in T221A mutant mice (Fig. 5c, 5d) and sleep rebound after SD was also reduced in T221A mutant mice (Fig. 5e), both supporting that T221 phosphorylation controls sleep need.

Our results that T221 phosphorylation enhances both the catalytic activity of SIK3 and the stability of SIK3 provide mechanistic basis to understand the function of T221 phosphorylation. It will be very interesting to study enzymes and mechanisms regulating T221 phosphorylation.

Our results with *Sik3-kof* mutants support that SIK3 is necessary for sleep need. Both sleep need measured by the δ power of EEG and sleep rebound after SD were decreased in *Sik3-kof* mice. By contrast, while GOF mutations in SIK1 and SIK2 increased sleep need ^29,34^, we report findings here that LOF mutations of SIK1 (Extended Data Fig. 1) or SIK2 (Extended Data Fig. 2) did not change sleep need, thus there is no evidence supporting that either SIK1 or SIK2 is physiologically necessary for sleep need in mice.

On the other hand, while the S551 site in SIK3 functionally important for sleep need ^27,29^, but MS measurement found no change in S551 phosphorylation after SD. This result did not support S551 as an indicator of sleep need.

Previous findings of increased sleep need with higher SIK3 activity in *Sleepy* and S551A mutants ^27,29,34^ can be re-interpreted to support the idea that SIK3 activity indicates and controls sleep need, complementing our results from *Sik3-kof* and SIK3 T221A mutants. It will be interesting to examine in *Sleepy* and S551A mutant mice whether sleep rebound after SD is increased with higher SIK3 activity in these mutant mice, which would further support a role for SIK3 activity in controlling sleep need. It will also be interesting to investigate whether T221 phosphorylation is increased in *Sleepy* and S551A mutant mice. A finding of increased T221 phosphorylation in *Sleepy* and S551A mutants will support that T221 is the key to monitor and control sleep need. If T221 phosphorylation is not increased in *Sleepy* and S551A mutants, it will suggest that S551 may regulate SIK3 activity in other manners or that it regulates other proteins downstream of SIK3 such as 14-3-3 to control sleep need ^42^.

Because T221 phosphorylation is gradual and dose-sensitive, it is not surprising that there is dosage sensitivity of SIK3 gene, mRNA and protein with regard to sleep need and sleep rebound after SD. The sleep phenotypes of +/*Sik3-kof* mice were sometimes similar to those of +/+ mice and other times similar to those of *Sik3-kof*/*Sik3-kof* mice (Fig. 4), paralleling the fact that the dosages of both SIK3 mRNA (Fig. 2b) and SIK3 protein (Fig. 2d) in +/*Sik3-kof* mice were in-between those of +/+ mice and *Sik3-kof*/*Sik3-kof* mice. Similarly, sleep phenotypes of +/T221A mice were sometimes similar to those of +/+ mice and other times similar to those of T221A/ T221A mice (Fig. 5), with the dosage of SIK3 protein in +/ T221A mice also in-between those of +/+ mice and T221A/ T221A mice (Fig. 2e).

SIKs were found more than two decades ago ^30-33^ and belong to adenosine monophosphate-activated protein kinase (AMPK) related kinases. Unlike AMPK, SIKs do not monitor cellular energy levels ^35^. SIKs play multiple physiological roles ^37^such as bone development ^47,48^, circadian rhythms ^49-51^, metabolism ^35,40^and sleep ^27,29,34,52^, and pathological roles ^36,37^such as inflammation ^53-55^ and cancer ^56-61^. It is curious whether other members of the AMPK related kinases play roles in sleep need or sleep.

Within their kinase domains, T182 in SIK1, T175 in SIK2 and T221 in SIK3 are equivalent to T172 in the catalytic α subunit of AMPK ^53-55,62^. T172 is important for AMPKα function but it is not known whether T172 phosphorylation in AMPKα is regulated by sleep or sleep need. It will be interesting to investigate potential regulation of AMPKα T172 phosphorylation by sleep, functional significance of AMPKα T172 phosphorylation in sleep and potential roles of sleep regulated AMPKα T172 phosphorylation in metabolic regulation.

Sleep is a crucial physiological process conserved in animals ^63^. There are two SIKs in *Drosophila* ^64^ and one in the worm *C. elegans* ^52,65^. It will be interesting to investigate their roles in sleep need, if reliable measure for sleep need can be established.

Molecular understanding of sleep regulation has heavily relied on the genetic approach, especially in the mouse and *Drosophila* by others and us over the last two decades ^25,66,67^. A breakthrough in understanding chemical changes associated with or responsible for regulating sleep need should stimulate further mechanistic research into sleep need and homeostatic regulation of sleep.

It has not escaped our notice that the chemical indication we have discovered immediately suggests a novel approach to study endogenous mechanisms and exogenous means that regulate sleep.

## Materials and Methods

### Antibodies

The primary antibodies used were anti-HDAC pS632(3424s, CST), anti-FLAG M2 HRP conjugated (A8592, Sigma), anti-SIK3 (Santa Cruz, sc-515408), anti-SIK3 pT221 (Abcam, ab271963), anti-PER2 (Abcam, ab227727), anti-PER2 pS662 (Abcam, ab206377), anti-Actin (Santa Cruz, sc-8342), anti-JNK (CST, 9252), anti-phospho-JNK (CST, 9251), anti-ERK1/2(CST, 4695), anti-phospho-ERK (CST, 4370).

### Cell Culture and Transfection

HEK293T cells were obtained from ATCC and cultured in Dulbecco’s modified Eagle’s medium (DMEM) medium containing 10% fetal bovine serum (FBS; Gibco) and 1% Penicillin/Streptomycin (Gibco). HEK293T cells were transfected with Lipofectamine 3000 reagent (Thermo Fisher) according to the manufacturer’s instructions and harvested 24 hrs-28hrs after transfection.

### Immunoprecipitation and Kinase Assay in vitro

Plasmids expressing FLAG-tagged human SIK3 and mutants were transfected into HEK293T cells for immunoprecipitation. After 24 hrs, cells were collected and lysed with 1 ml lysis buffer (25 mM Tris-HCl pH7.5, 150 mM NaCl,5 mM MgCl2, 0.5% NP40, 0.25 M Sucrose, 1 mM DTT, 1 mM PMSF, 1x protease inhibitor cocktail (Roche), 1x phosphatase inhibitor II,1x phosphatase inhibitor III (Sigma), before centrifugation of cell lysates at 20000 g for 30 min at 4 □. Cell lysates were incubated with 10μl anti-FLAG antibody coated magnetic agarose beads, balanced by lysis buffer for 1 hr. Agarose beads were washed with 1 ml lysis buffer three times at 4 □, before final washing with 100 μl 0.5mg/ml 3xFLAG peptide buffer A at room temperature and storage at -80 □. Kinase reactions were performed for 30 mins at 37 □ by adding 10 μl kinase, 1 μg HDAC4 with a final concentration of 1 mM ATP. Samples were analyzed by immunoblotting with the indicated antibodies.

### Expression and Purification of Recombinant Proteins

cDNAs for specific genes were cloned into pET-28a vector with indicated tags. Plasmids were transformed into *E. coli* BL21 cells and induced with 0.5 mM isopropyl-β-D-thiogalactoside (IPTG) at 18 □ for 16 hrs. Cells were harvested and suspended in Ni binding buffer (300 mM NaCl, 20 mM Tris-HCl, pH 7.5) supplemented with protease inhibitors. Cells were lysed by sonication and centrifuged at 14000 rpm for 30 min. Supernatants were filtered by 0.45 μm filter and purified to 90% purity by tandem Ni^2+^ and dextran affinity column. Pooled elution fractions were stained by Coomassie blue and stored at -80□.

### Animals

*Sik3 knockout first(sik3-kof)* mice were purchased from CAM-SU GRC (Clone No. HEPD0744_6_E02). *Sik1 knockout (Sik1ko), Sik2 knockout (Sik2ko)* and *Sik3* point mutation mice were made as described later.

All experimental procedures were performed in accordance with the guidelines and were approved by the IACUC of Chinese Institute for Brain Research, Beijing. Mutant mice and wt littermates were maintained on a C57BL/6N background. Mice were housed under a 12 hr:12 hr light/dark cycle and controlled temperature and humidity conditions. Food and water were delivered *ad libitum*.

### Mouse Protein Preparation

Whole brains of mice were quickly dissected, rinsed with 1 X PBS and homogenized by homogenizer (Wiggens, D-500 Pro) in ice-cold lysis buffer (20 mM HEPES, 10 mM KCL, 1.5 mM MgCl2, 1 mM EDTA, 1 mM EGTA, 1 mM DTT, freshly supplemented with a protease and phosphatase inhibitor cocktail). Brain homogenates were centrifuged at 14000 rpm for 15 min. Supernatants were carefully transferred to a new microtube. Protein concentrations of brain lysates were determined with the bicinchoninic acid (Thermo Fisher, 23225) assay and normalized to 2 mg/ml. Before immunoblotting, samples were kept in liquid nitrogen, if necessary.

### Quantitative Real-Time PCR

RNA was extracted from mouse brain lysates (mixture of 3∼4 mouse brains for each sample) using RNAprep pure Tissue Kit (Tiangen, DP431). cDNA was synthesized using HiScript II 1st Strand cDNA Synthesis Kit (Vazyme, R211-01). Quantitative real-time PCR were performed using ChamQ SYBR qPCR Master Mix (Vazyme, R211-01) in the Roche LightCycler® 96 Instrument. The following primers were used:

Sik1-Fw (5’-TCTGTCCTGCAGGCTGAGAT-3’),

Sik1-Rv (5’-ATAGCTGTGTCCAGCAGACT-3’),

Sik2-Fw (5’-TGAGCAGGTTCTTCGACTGAT-3’),

Sik2-Rv (5’-AGATCGCATCAGTCTCACGTT-3’),

Sik3-Fw (5’-TTGTCAATGAGGAGGCACAC-3’),

Sik3-Rv (5’-TGTTAGGGGCCACTTGTAGG-3’),

Gapdh-Fw (5’-AGAACATCATCCCTGCATCC-3’),

Gapdh-Rv (5’-CACATTGGGGGTAGGAACAC-3’).

### Generation of point mutation and knock out mice

To select proper single-guide RNA (sgRNA), candidate sgRNAs were inserted into pCS-3G vector through Gibson assembly method and the cleavage activity was verified by UCA™ method (Biocytogen). For each mutant mice, two sgRNAs were selected and transcribed in vitro to obtain RNAs (*Sik1ko*: 5’-gcgggttgaggacgcacgggtgg-3’ and 5’-gccagaagaccgccttgtcgtgg-3’; *Sik2ko*: 5’-agtctgtctcgcaatggaccagg-3’ and 5’-tcagtgtagctccagatatgtgg-3’; *Sik3ko*: 5’-agtcattaacaacgactatgcgg-3’ and 5’-gtccctcgcaacccaagcgccgg-3’ and *Sik3*^*221A/A*^: 5′-aggcgacatacctacctccctgg-3′ and 5′-cgcggttggcactgatgcccagg-3′). For *Sik3*^*221A/A*^, an additional donor vector was constructed with two ∼1.5kb homologous arms and donor sequence which covered exon 5 and 6 with nucleotide mutation from ACG to GCG(T221A) and additional sequence mutation to introduce AseI and BgIII restriction enzyme sites for identification. sgRNAs only or sgRNAs and donor vector were injected into C57BL/6N mouse embryos. F0 mice were genotyped for the presence of the recombination. F0 mice were mated with the counter C57BL/6N mice to obtain F1 offspring. The successful recombination was identified through Southern blot analysis in F1 mice. Mutant lines were back-crossed to C57BL/6N for at least 5 generations to exclude possible off-targeting.

### Electrode Implantation Surgery

EEG & EMG electrodes were made by sealing two EEG screws and two EMG wires in a 2*2 pin header with stainless steels (COONER WIRE™ AS633). The underside of pin header was covered by silicone rubber (Ausbond^R^ 704) for insulation. Self-made electrodes were implanted into mice under anesthesia using isoflurane (3% for induction, 1.5% for maintenance) for EEG (AP-1mm/ML-1.5mm relative to bregma, AP-1mm/ML-1.5mm relative to posterior fontanelle) and EMG (two wires inserted in the left and right neck muscles separately) acquisition. After implantation, the pin header was attached to the skull first by glue and then by dental cement. All mice were maintained individually for at least 5 days to recover from surgery. After recovery, mice were placed in the recording cage and tethered to an omni-directional arm (RWD Life Science Inc.) with connection cable for 2days of habituation before recording.

### Sleep Recording and Analysis

EEG and EMG data were recorded at a sampling frequency of 200 Hz with custom software (based on LabView 2020, National Instruments Corp.) for 2 consecutive days to analyze sleep/wake behavior under baseline conditions. The recording room was kept under a 12 hr:12 hr light/dark cycle and a controlled temperature (24-25□). EEG/EMG data were initially processed by Accusleep^68^, and then manually checked in SleepSign™ to improve accuracy and reduce analysis time. The data was classified in a 4s epoch, the sleep stage was classified as reference^67^. Briefly, wakefulness was scored as high amplitude and variable EMG and fast and low amplitude EEG; NREM was scored as high amplitude δ (1-4 Hz) frequency EEG and low EMG tonus; REM was characterized as complete silence of EMG signals and low amplitude high frequency θ(6-9 Hz)-dominant EEG signal.

For power spectrum analysis, EEG was subjected to fast Fourier transform (FFT) analysis with a hamming window method by SleepSign™, yielding power spectra between 0-25 Hz with a 0.39Hz bin resolution. State EEG power spectra represent the mean of power distribution for each state-corresponding epoch, i.e. NREMS, REMS, WAKE, during a diurnal recording period. The hourly NREMS δ-power density indicates the average of δ-power density as a percentage of δ-band power (1-4 Hz) to total power (0-25 Hz) for each NREM epoch contained in an hour. Epochs containing movement artifacts were marked and excluded automatically in the spectra analysis by a custom algorithm, which initialy summed up the power for each epoch at all frequency bins, calculating the difference of total power between each adjacent epochs and figuring out the abnormal value among these differences. The threshold T of differences was determined by visual inspection based on the distribution graph of these differences, epochs which contributed to those differences greater than T were identified and discarded. The state episode was defined as a continuous and single state epoch, NREM episode was defined as at least 3 continuous NREM epochs and was ended by REM epoch or at least 3 wakefulness epochs. REM or WAKE episode consists of at least 2 or 3 consecutive REM epochs or wakefulness epochs, respectively. The number and average duration of episode were defined as episode number and episode duration, respectively.

### Sleep Deprivation

Sleep deprivation was manually carried out by preventing mice from sleep using a stick during ZT 0-6. This will ensure mice were awake during deprivation and reduce additional stress. Rebound of each stage was calculated as cumulative changes of WAKE/NREM/REM duration after SD compared to duration in the same ZT before SD. Changes of δ power was calculated as δpower post-SD minus δ power pre-SD.

### Statistics

Statistical analysis was performed with Graph Pad Prism 8.0.2 (GraphPad Software).

Kruskal-Wallis test followed by Dunn’s posttest was used to compare mean results from different lines. Two-way ANOVA followed by Turkey’s multiple comparisons test was used to compare power spectrum between different lines. Statistical significance is denoted by asterisks: \**P < 0*.*05*, \*\**P < 0*.*01*, \*\*\**P < 0*.*001*. Error bars in the graphs represent mean ± SEM.

## SUPPLEMENTAL INFORMATION

Supplemental information includes 4 figures.

## ACKNOWLEDGEMENTS

We are grateful to the Beijing Commission of Science and Technology for grant support.

## Legends for Extended Data Figures

**Extended Data Figure 1.**
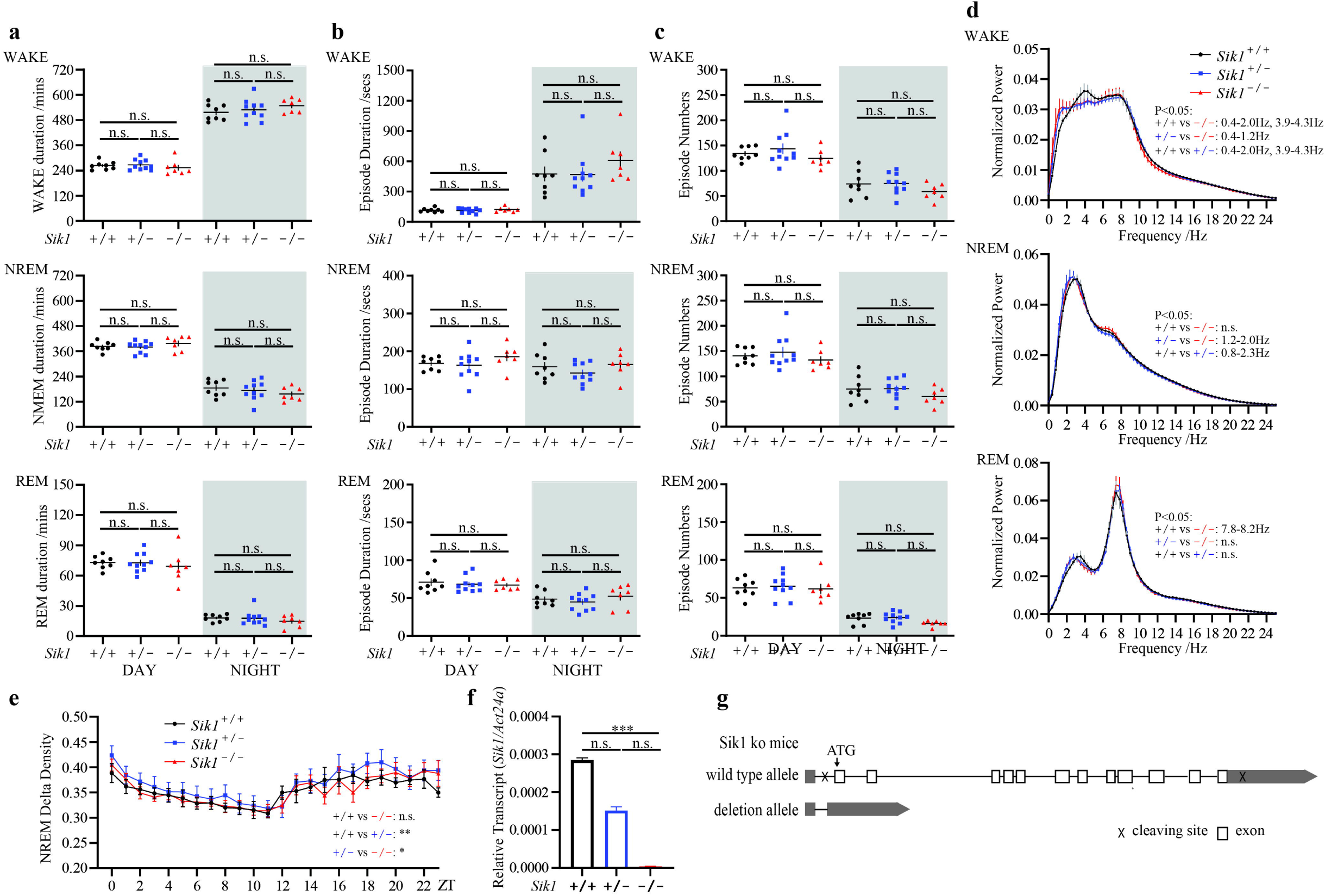
*Sik1* and Sleep in Mice. **a**. Duration of WAKE, NREM and REM in *Sik1*^*+/+*^(n=8), *Sik1*^*+/-*^(n=10) and *Sik1*^*-/-*^(n=7) mice. Kruskal–Wallis test with Dunn’s posttest. Error bars represent SEM. **b**. Episode duration of WAKE, NREM and REM in *Sik1*^*+/+*^(n=8), *Sik1*^*+/-*^(n=10) and *Sik1*^*-/-*^(n=7) mice. Kruskal–Wallis test with Dunn’s posttest. Error bars represent SEM. **c**. Episode number of WAKE, NREM and REM in *Sik1*^*+/+*^(n=8), *Sik1*^*+/-*^(n=10) and *Sik1*^*-/-*^(n=7) mice. Kruskal–Wallis test with Dunn’s posttest. Error bars represent SEM. **d**. EEG power spectra of WAKE, NREM and REM states in *Sik1*^*+/+*^(n=8), *Sik1*^*+/-*^(n=10) and *Sik1*^*-/-*^(n=7) mice. Two-way ANOVA with Tukey’s posttest, *P*<0.05 pairs were listed on the right. Error bars represent SEM. **e**. NREM δ-power density of *Sik1*^*+/+*^(n=8), *Sik1*^*+/-*^(n=10) and *Sik1*^*-/-*^(n=7) mice. Two-way ANOVA with Tukey’s posttest. \**P*< 0.05, \*\**P*<0.01. Error bars represent SEM. **f**. Relative *Sik1* mRNA expression levels in the brains of *Sik1*^*+/+*^(n=4), *Sik1*^*+/-*^(n=6) and *Sik1*^*-/-*^(n=6) mice. Kruskal–Wallis test with Dunn’s posttest. \*\*\**P*<0.001. Error bars represent SEM. **g**. Schematic representations of mouse *Sik1* gene and the region deleted from *Sik1*^*-/-*^ mutants. Asingle transcript (NM_010831.3) generates a protein of 779 aa (NP_034961.2) in wt and the entire coding region was deleted in *Sik1*^*-/-*^ mutant.

**Extended Data Figure 2.**
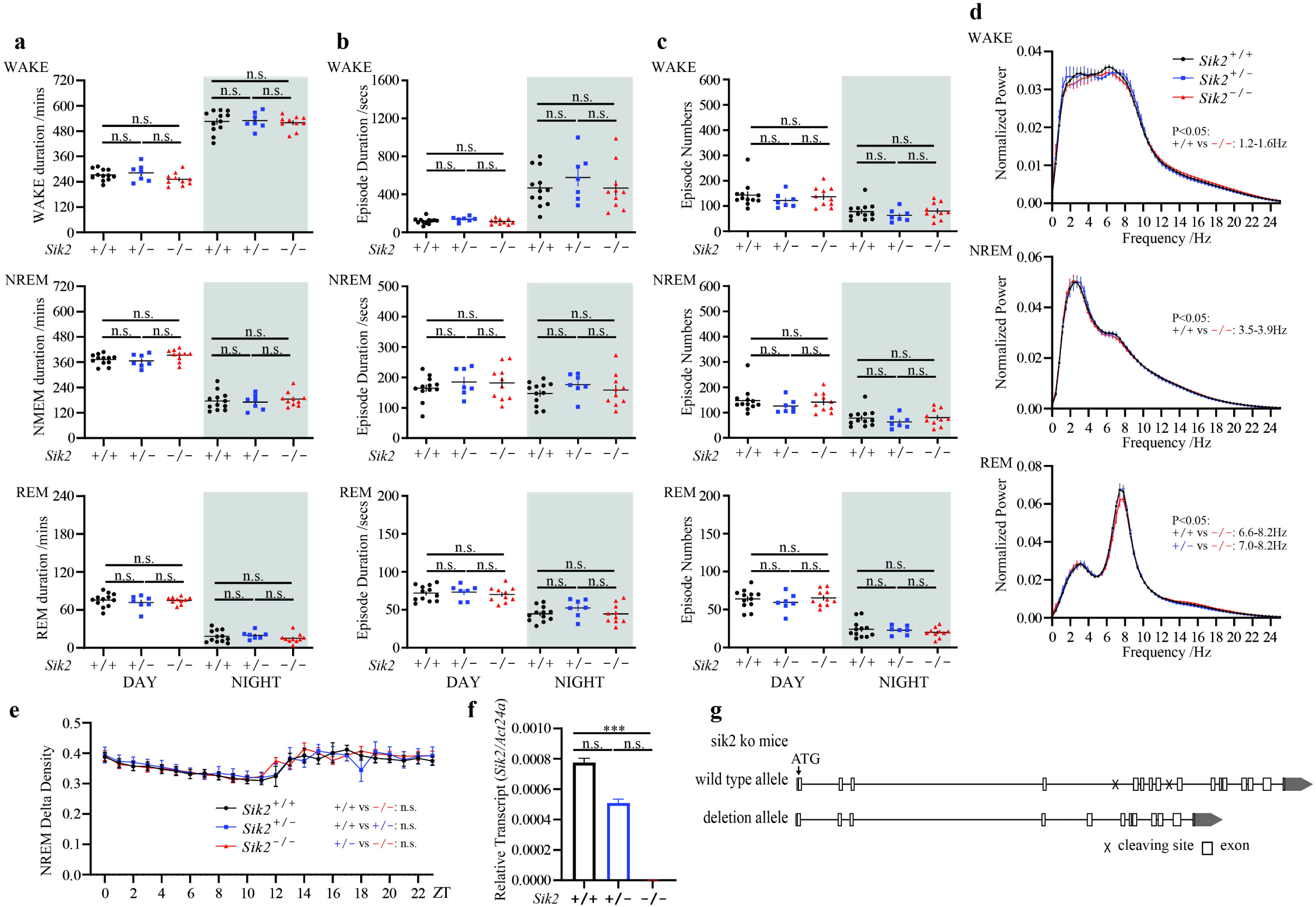
*Sik2* and Sleep in Mice. **a**. Duration of WAKE, NREM and REM in *Sik2*^*+/+*^(n=12), *Sik2*^*+/-*^(n=7) and *Sik2*^*-/-*^(n=10) mice. Kruskal–Wallis test with Dunn’s posttest. Error bars represent SEM. **b**. Episode duration of WAKE, NREM and REM in *Sik2*^*+/+*^(n=12), *Sik2*^*+/-*^(n=7) and *Sik2*^*-/-*^ (n=10) mice. Kruskal–Wallis test with Dunn’s posttest. Error bars represent SEM. **c**. Episode number of WAKE, NREM and REM in *Sik2*^*+/+*^(n=12), *Sik2*^*+/-*^(n=7) and *Sik2*^*-/-*^(n=10) mice. Kruskal–Wallis test with Dunn’s posttest. Error bars represent SEM. **d**. EEG power spectra of WAKE, NREM and REM states in *Sik2*^*+/+*^(n=11), *Sik2*^*+/-*^(n=7) and *Sik2*^*-/-*^ (n=10) mice. Two-way ANOVA with Tukey’s posttest, *P*<0.05 pairs were listed on right. Error bars represent SEM. **e**. NREM δ-power densities of *Sik2*^*+/+*^(n=11), *Sik2*^*+/-*^(n=7) and *Sik2*^*-/-*^(n=10) mice. Two-way ANOVA with Tukey’s posttest. Error bars represent SEM. **f**. Relative *Sik2* mRNA expression levels in the brains of *Sik2*^*+/+*^(n=6), *Sik2*^*+/-*^(n=6) and *Sik2*^*-/-*^ (n=6) mice. Kruskal–Wallis test with Dunn’s posttest. \*\*\**P*<0.001. Error bars represent SEM. **g**. Schematic representations of mouse *Sik2* gene and the region deleted in *Sik2*^*-/-*^ mutant. Asingle transcript (NM_178710.3) generates a protein of 931aa (NP_848825.2) in wt and exon5-exon8 was deleted in *Sik2*^*-/-*^ mutant resulting in the deletion of aa160-aa367 and a frame shift.

**Extended Data Figure 3.**
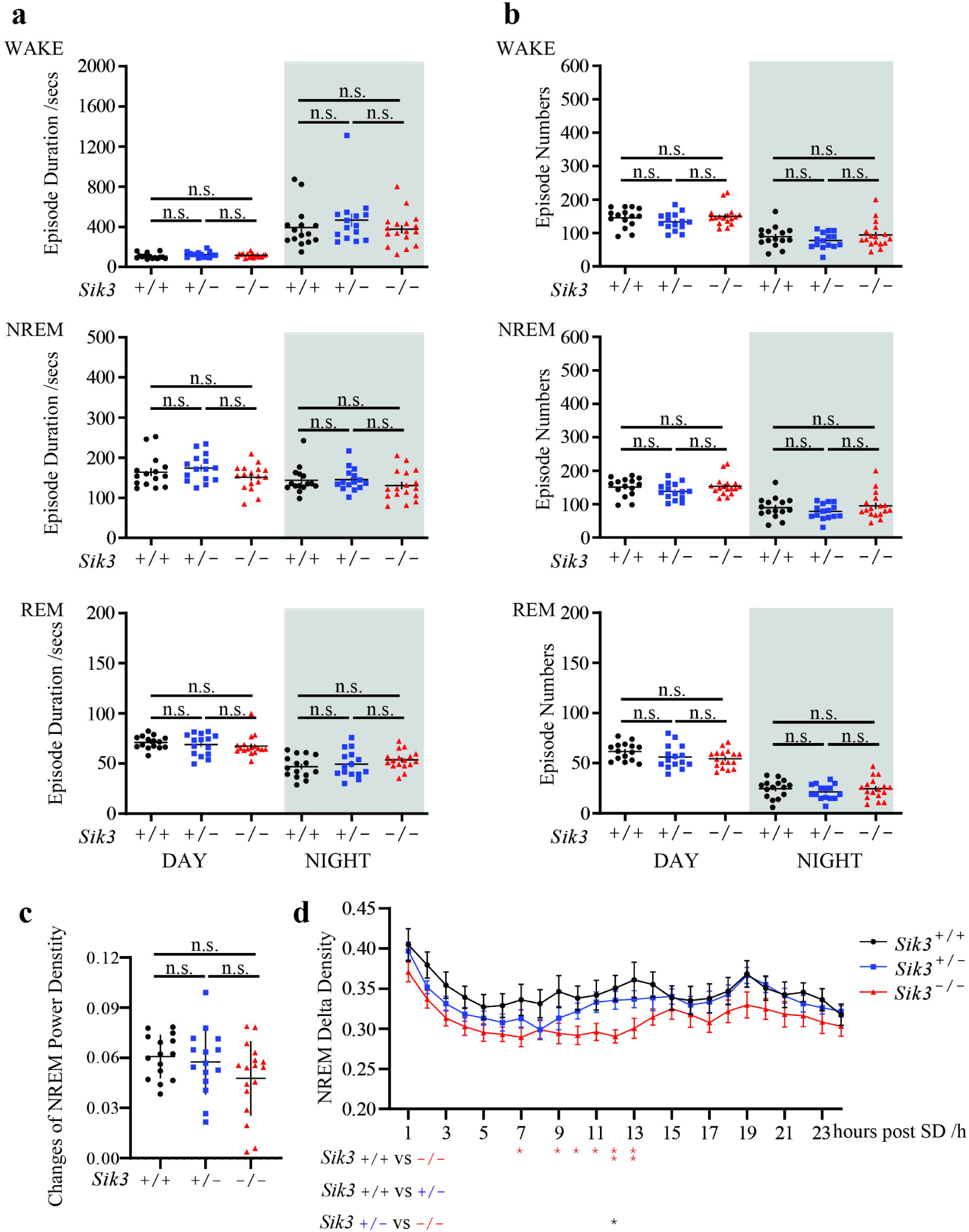
Sleep Need *Sik3-kof* Mutants after SD. **a**. Episode duration of WAKE, NREM and REM in *Sik3*^*+/+*^(n=15), *Sik3*^*+/-*^(n=15) and *Sik3*^*-/-*^ (n=17) mice. Kruskal–Wallis test with Dunn’s posttest. Error bars represent SEM. **b**. Episode number of WAKE, NREM and REM in *Sik3*^*+/+*^(n=15), *Sik3*^*+/-*^(n=15) and *Sik3*^*-/-*^ (n=17) mice. Kruskal–Wallis test with Dunn’s posttest. Error bars represent SEM. **c**. δ power density increasement from ZT9-11 to ZT19-24 of *Sik3*^*+/+*^(n=15), *Sik3*^*+/-*^(n=15) and *Sik3*^*-/-*^(n=17) mice. Kruskal–Wallis test with Dunn’s posttest. Error bars represent SEM. **d**. NREM δ-power densities after 6h sleep deprivation of *Sik3*^*+/+*^(n=11), *Sik3*^*+/-*^(n=12) and *Sik3*^*-/-*^ (n=11) mice. Two-way ANOVA with Tukey’s posttest, comparation results between genotypes were listed below. \**P*< 0.05, \*\**P*<0.01. Error bars represent SEM.

**Extended Data Figure 4.**
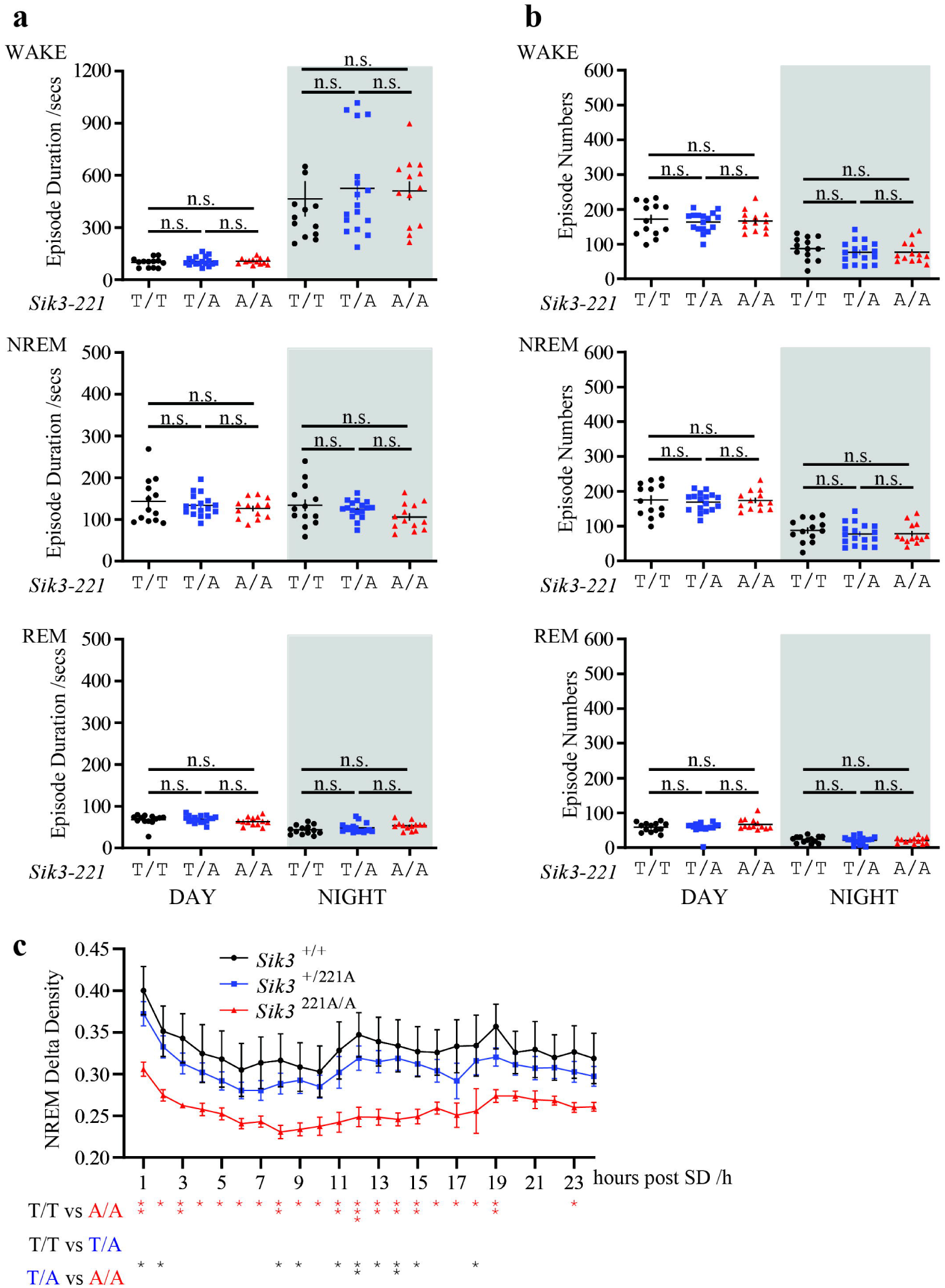
Sleep Need of *Sik3-T221A* Mutants after SD. **a**. Episode duration of WAKE, NREM and REM in *Sik3*^*+/+*^(n=13), *Sik3*^*+/221A*^(n=17) and *Sik3*^*221A/A*^ (n=13) mice. Kruskal–Wallis test with Dunn’s posttest. Error bars represent SEM. **b**. Episode number of WAKE, NREM and REM in *Sik3*^*+/+*^(n=13), *Sik3*^*+/221A*^(n=17) and *Sik3*^*221A/A*^ (n=13) mice. Kruskal–Wallis test with Dunn’s posttest. Error bars represent SEM. **c**. NREM δ-power density after 6 hr SD of *Sik3*^*+/+*^(n=6), *Sik3*^*+/221A*^(n=10) and *Sik3*^*221A/A*^ (n=8) mice. Two-way ANOVA with Tukey’s posttest, comparation results between genotypes were listed below. \**P*<0.05, \*\**P*<0.01, \*\*\**P*<0.001. Error bars represent SEM.

